# Dibenzoylmethane, a novel β-diketone pore blocker of large-conductance calcium-activated potassium channel

**DOI:** 10.64898/2026.04.24.720584

**Authors:** Piotr Koprowski, Przemysław Miszta, Jakub W. Strawa, Yurii Krempovych, Alicja Ziajowska, Sławomir Filipek, Adam Szewczyk, Michał Tomczyk

**Author notes:** P.K. and P.M. contributed equally to the work. Corresponding author. Laboratory of Intracellular Ion Channels, Nencki Institute of Experimental Biology, Polish Academy of Sciences, ul. Pasteur 3, 02-093 Warsaw, Poland., *E-mail address* (Piotr Koprowski).

## Abstract

Large-conductance calcium-activated potassium (BK_Ca_) channels are ubiquitously expressed in mammalian cells and regulate electrical activity, intracellular calcium signaling, and cell survival. Although BK_Ca_ dysfunction has been linked to multiple diseases, the number of selective channel modulators is limited. In this study, we characterize dibenzoylmethane (DBM), a plant-derived compound isolated from *Hottonia palustris*, as a novel inhibitor of BK_Ca_ channel activity in both plasma membrane and mitochondrial BK_Ca_. Electrophysiological recordings revealed that DBM lowers the open probability of BK_Ca_ channels in a concentration-dependent fashion and markedly reduces mean open time, leading to a pronounced flickering behavior - hallmarks of pore-targeted blockade. Competition experiments demonstrated that DBM antagonizes the effect of paxilline, a high-affinity pore-binding inhibitor, suggesting overlapping binding sites. Molecular dynamics simulations further supported this hypothesis, showing that several DBM molecules can block the pore by employing π-π interactions with each other and pore residues. On top of the pore, the carbonyl groups of DBM block the nearest potassium ion in the selectivity filter. The presence of DBM induces the removal of water molecules from the pore. To assess the structural requirements for activity, we tested three DBM analogs: phenyl-1,3-butanedione (PBD), *trans*-chalcone (T-Ch), and (E)-1,3-diphenylprop-2-en-1-ol (DPE). T-Ch and DPE inhibited BK_Ca_ channels with comparable efficacy to DBM, whereas PBD was significantly less potent. These results indicate that diphenyl substitution and structural rigidity are critical determinants of inhibitory activity. Our findings position DBM and its analogs as promising chemical scaffolds for the development of selective BK_Ca_ channel modulators with potential pharmacological applications.

## 1. Introduction

Large-conductance calcium- and voltage-activated potassium (BK_Ca_, K_Ca_1.1) channels couple membrane depolarization to intracellular Ca^2+^ signals and provide powerful negative feedback control of excitability, secretion, and smooth muscle tone due to their unusually high single-channel conductance. BK_Ca_ channels are formed by tetramers of the pore-forming Slo1 α subunit (KCNMA1) and may assemble with auxiliary β and γ subunits that tune gating, pharmacology, and trafficking in a tissue- and context-dependent manner (Gonzalez-Perez and Lingle, 2019). Recent cryo-EM structures of human Slo1-containing complexes have resolved key features of the pore-gate domain and central cavity that are critical for interpreting and rationalizing pore-directed ligand interactions (Tao and Mackinnon, 2019). Natural products remain a major source of pharmacologically active small molecules, including modulators of K^+^ channels. Several flavonoids have been reported to activate BK_Ca_ channels in cellular systems, including quercetin and naringenin. Conversely, fungal indole-diterpenoids such as penitrem A, paxilline (and related compounds) are widely used BK_Ca_ inhibitors and are valuable tools for mechanism-based classification of new ligands (Asano et al., 2012; Zhou and Lingle, 2014). Collectively, these observations motivate continued screening of chemically diverse plant-derived scaffolds for BK_Ca_ channel modulation, particularly for ligands that act through pore-directed mechanisms.

BK_Ca_ channels are also functionally present in mitochondria (mitoBK_Ca_) (Bednarczyk et al., 2013; Gałecka et al., 2021; Singh et al., 2013; Szabo and Szewczyk, 2023), where they have been implicated in the regulation of mitochondrial bioenergetics, redox signaling, and susceptibility to stress pathways in a cell-type–dependent manner (Augustynek et al., 2018; Balderas et al., 2015; Kaczara et al., 2015; Kulawiak et al., 2023; Maliszewska-Olejniczak et al., 2024; Szewczyk, 2024; Walewska et al., 2022). Notably, BK-type activity in the inner mitochondrial membrane was initially demonstrated by patch-clamp recordings from mitoplasts derived from a human glioma cell line, establishing a link between mitoBK_Ca_ function and glial tumor models (Bischof et al., 2024; Siemen et al., 1999). In parallel, BK_Ca_ channel splice diversity and glioma-associated BK_Ca_ variants (“gBK”) have been described in glioma and glioblastoma samples and cell lines, supporting the broader concept that BK_Ca_ channel molecular heterogeneity may be relevant to tumor cell physiology (Ge et al., 2012; Liu et al., 2002). Consistent with this, BK_Ca_ channels have been repeatedly discussed as potential contributors to glioblastoma cell migration and invasion in specific experimental contexts (Catacuzzeno et al., 2015). Interestingly, inhibition of mitochondrial potassium channels such as Kv1.3 may lead to cell death (Szabo and Szewczyk, 2023).

Mechanistically, pore-directed BK_Ca_ inhibitors like quaternary amonium ions are particularly informative because they can generate characteristic single-channel signatures and show interpretable relationships with the gating state (Li and Aldrich, 2004; Tang et al., 2009; Wilkens and Aldrich, 2006). Paxilline (PAX), a canonical high-affinity BK_Ca_ inhibitor, has been shown to inhibit BK_Ca_ channels predominantly *via* a closed-state–dependent mechanism, with inhibition inversely related to channel open probability (Zhou and Lingle, 2014). Structure-guided and computational analyses further define determinants of PAX inhibition within the pore-gate domain and central cavity, providing a reference framework for interpreting novel pore-acting ligands (Zhou et al., 2020).

*Hottonia palustris* L. (Primulaceae) is a widely distributed aquatic plant reported to contain multiple bioactive metabolites, including dibenzoylmethane (DBM) (Strawa et al., 2022a, 2022b). Here, we investigate whether DBM modulates BK_Ca_ channels both in the plasma membrane and mitochondria. We define its mechanism of action using electrophysiology combined with molecular modeling. Using U-87MG astrocytoma cells as a model system, we test whether DBM produces functional signatures consistent with pore-directed blockade and assess structural hypotheses for DBM occupancy within the BK_Ca_ pore that may inform future β-diketone–based optimization. To assess the structural requirements for activity, we additionally tested three other DBM analogs: phenyl-1,3-butanedione (PBD), *trans*-chalcone (T-Ch), and (E)-1,3-diphenylprop-2-en-1-ol (DPE).

## 2. Methods

### 2.1. Cell culture

U-87MG astrocytoma cells were maintained in DMEM (General Chemistry Laboratory, Institute of Immunology and Experimental Therapy, Polish Academy of Sciences, Wroclaw, Poland) supplemented with 2 mM L-glutamine (Gibco, Carlsbad, CA, USA), 10% FBS (Gibco, Carlsbad, CA, USA), 100 U/mL penicillin (Sigma-Aldrich, St. Louis, MO, USA), and 100 µg/mL streptomycin (Sigma-Aldrich, St. Louis, MO, USA). The cells were incubated at 37°C in a humidified 5% CO₂ environment and were typically fed and reseeded every four days.

### 2.2. Chemicals

The dibenzoylmethane-containing fraction from *H. palustris* was prepared following the procedure outlined in our previous study (Strawa et al., 2022a). Dibenzoylmethane (DBM, 1,3-diphenylpropane-1,3-dione) was obtained in two polymorphic forms, and its structure and purity were confirmed by crystallographic and spectral analyses (NMR, MS) (Strawa et al., 2022b). The following small molecules were used in the study: 1-phenyl-1,3-butanedione (PBD) (99%, CAS No. 93-91-4, B11907, Merck), *trans*-chalcone (T-Ch) (98.72%, CAS No. 614-47-1, A131628, AmBeed), and (E)-1,3-diphenylprop-2-en-1-ol (DPE) (95%, CAS No. 62668-02-4, A260528, AmBeed), synthetic dibenzoylmethane (98%, CAS No.120-46-7, D33454, Merck), and paxilline (PAX) (98%, CAS No. 57186-25-1, P2928, Merck).

### 2.3. Mitochondria isolation

Mitochondria from U-87MG cells were isolated following established protocols with slight modifications. In brief, cells were rinsed with PBS (General Chemistry Laboratory, Institute of Immunology and Experimental Therapy, Polish Academy of Sciences, Wroclaw, Poland), scraped, suspended in PBS, and subjected to centrifugation at 800× g for 10 minutes. The cell pellet was resuspended in isolation buffer (250 mM sucrose, 1 mM EGTA, 5 mM HEPES, pH 7.2) and homogenized. The homogenate was centrifuged at 9200× g for 10 minutes in an Eppendorf tube. After resuspending the pellet in isolation buffer, it was centrifuged at 780× g for 10 minutes, and the supernatant containing mitochondria was transferred to a new tube for a final centrifugation at 9200× g for 10 minutes. The resulting mitochondrial pellet was resuspended in a small volume of isolation buffer (20–100 µL). All procedures were performed on ice, and all centrifugations were conducted at 4°C. Isolated mitochondria were kept on ice and used for patch-clamp experiments within 4 to 6 hours of isolation.

### 2.4. Mitoplast preparation

Mitoplasts (inner mitochondrial membranes) were prepared using a previously established method. Briefly, 1–2 µL of isolated mitochondria from U-87MG cells were incubated in 40 µL of hypotonic solution (5 mM HEPES, 100 µM CaCl₂, pH 7.2) for approximately 2 minutes to induce swelling and outer membrane rupture. After this, 10 µL of hypertonic solution (1.5 M sucrose, 30 mM HEPES, 100 µM CaCl_2_, pH 7.2) was added to restore isotonicity.

### 2.5. Patch-clamp experiments

Patch-clamp studies were conducted as described in previous publications. Experiments were performed in the inside-out configuration. Freshly scraped cells or mitoplasts (0.5–2 µL) were introduced directly into the recording chamber containing a high calcium solution (150 mM KCl, 10 mM HEPES, and 100 µM CaCl_2_, pH 7.2). Channel activity modulators were applied directly into the chamber (for both inside-out and outside-out configurations). Patch-clamp pipettes, fabricated from borosilicate glass (Harvard Apparatus GC150–10, Holliston, MA, USA) with a resistance of ∼15 MΩ, were filled with high calcium solution. Upon capturing the mitoplast and forming a giga-ohm seal, the membrane patch was excised by lightly tapping the pipette holder. Ionic currents were recorded using a patch-clamp amplifier (Axopatch 200B, Molecular Devices Corporation, San Jose, CA, USA), low-pass filtered at 1 kHz, and sampled at 2.5 kHz. The probability of mitoBK_Ca_ channel opening (P(o)) was calculated using Clampfit 10.7 software’s single-channel search mode. The number of stars in the figures represents statistical significance between two data sets: * p < 0.05, ** p < 0.01, *** p < 0.001. “n” shown in captions of figures represents the number of patches used in analysis. Normalisation and statistical analysis were performed using GraphPad Prism 10. Concentration–response relationships shown in Fig. 2C were fitted in GraphPad Prism 10 using the Hill equation:

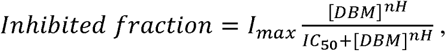

with [DBM] □ dibenzoylmethane concentration, I_max_ □ maximal inhibition, IC_50_ □ concentration producing half-maximal inhibition, n_H_ □ Hill coefficient. This provided the best or near-best fit for the datasets recorded at positive membrane potentials. For datasets recorded at negative membrane potentials, fitting was less reliable because channel inhibition was already close to maximal across most of the tested concentration range.

### 2.6. Molecular modelling

#### 2.6.1. Protein preparation

The initial structure of the human BK_Ca_ channel in open form was obtained from the Protein Data Bank (PDB ID: 6V38) (Tao and Mackinnon, 2019). The structure was determined by cryo-electron microscopy at a resolution of 3.8 Å in the presence of calcium ions, which activated the channel. Only the transmembrane domain of the tetrameric BK_Ca_ channel was retained for further simulations, as this region contains the pore that can be blocked by ligands and preclude the ion permeation. All non-essential molecules, such as cholesterol and stabilizing agents used during structure determination, were removed, and the modified residues were converted to standard ones. Hydrogen atoms were added to generate residue protonation states corresponding to physiological pH. Due to missing residues (52–92) in the long and flexible loop between helices S0 and S1 and the absence of the first 18 N-terminal residues, the entire helix S0 was removed from each of the four subunits of BK_Ca_. Helix S0 has only a small contact with the rest of the channel, so its stabilizing role is negligible. It is a first helix in a voltage-sensing domain (VSD) composed of S0–S4 helices, but since we studied ligand docking to the pore-gating domain (PGD), involving inner transmembrane helices S5 and S6, the omission of S0 in ligand docking and subsequent MD simulations is justified. To study potassium ion permeability, we added four potassium ions to the selectivity filter since they were not present in the 6V38 structure. The ions were copied to the 6V38 structure from the BK_Ca_ channel of *Aplysia californica* (the California sea hare) (PDB ID: 5TJ6) (Tao et al., 2017) after structural alignment of both structures.

#### 2.6.2. Membrane embedding

A membrane bilayer composed of pure POPC (1-palmitoyl-2-oleoyl-*sn*-glycero-3-phosphocholine) was generated using the Membrane Builder extension in VMD 1.9.3 (Humphrey et al., 1996). The bilayer was created using 412 POPC molecules, resulting in a square membrane with dimensions 107.5 Å × 107.5 Å. This was the smallest box to fit the whole BK_Ca_ channel and keep a 15 Å distance from the membrane edges. To make the best fit of BK_Ca_ to the membrane, the VSD domains of BK_Ca_ were located in the corners of the membrane square. The protein orientation in the membrane was set according to the Orientations of Proteins in Membranes (OPM) database (Lomize et al., 2012). The lipids overlapping the protein were removed, and the final membrane contained 232 lipids. The system was subsequently solvated with TIP3P water layers on both sides of the membrane, resulting in a final box dimension of 107.5 Å × 107.5 Å × 117.8 Å.

#### 2.6.3. Ligand preparation

Dibenzoylmethane (DBM) structure was retrieved from the PubChem database (Kim et al., 2025) and prepared in the LigPrep module of the Schrödinger suite of programs (v. 2024-2). LigPrep, using the OPLS4 force field (Lu et al., 2021), generated five keto-enol tautomers, including ionized states, existing at target pH 7.0 ± 2.0 (Fig. 1A). All forms were subjected to 200 ns MD simulations in water environment, and the diketo form proved to be the most flexible due to the absence of the C=C double bond. Since the ligand’s flexibility is important to fit the pore region, this form was selected for the ligand docking.

**Fig. 1.**
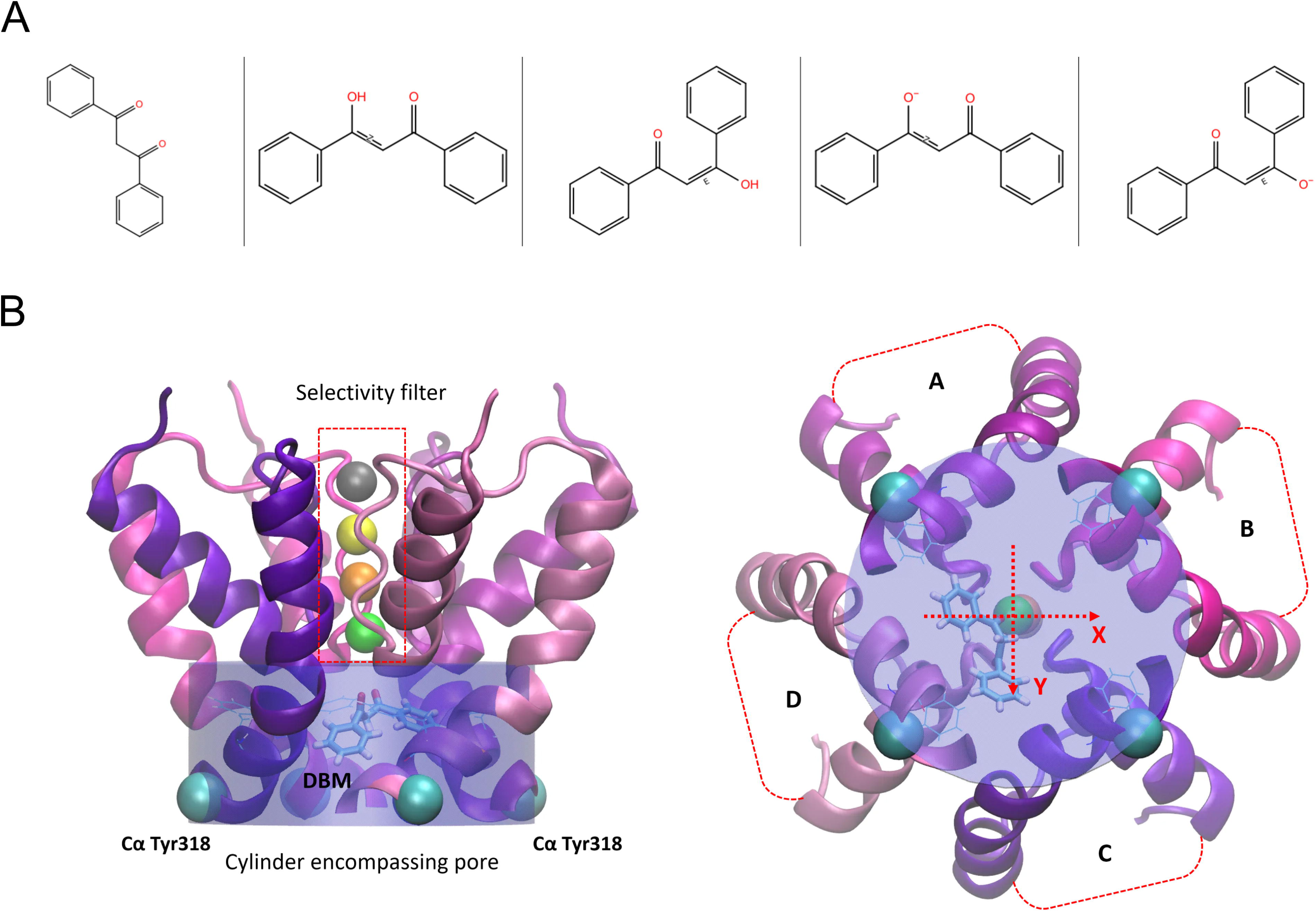
Docking setup for dibenzoylmethane (DBM) in the BK_Ca_ channel pore. (A) Structures of dibenzoylmethane (DBM) tauto-conformers generated at neutral pH (7.0 ± 2.0) and considered during ligand preparation for molecular docking. The diketo form shown on the left was used for docking and subsequent model preparation. (B) Visualization of the cylindrical region used for analysis of pore hydration and cavity volume during molecular dynamics simulations. Side view (left) and cytoplasmic view (right) are shown. The coordinate axes used to track DBM movement are centered in the middle of the cylinder.

#### 2.6.4. Ligand docking and the system completion

The DBM molecules were docked to the channel pore using AutoDockVina v1.1.2 software, which enabled docking of several ligand copies at once. The ligand and protein structures were prepared using the DockPrep tool (DOCK 6.11) that comes with UCSF Chimera 1.17.3. Partial atomic charges for standard residues were obtained from AMBER ff14SB (Maier et al., 2015), while charges for nonstandard residues were calculated using an Antechamber module in UCSF Chimera. The search volume for ligand docking in the BK_Ca_ channel was set to include the entire pore region. We docked one, two, and four DBM molecules to the pore region. In each case, the poses with the highest docking score were selected. As many as four ligands were able to fit into the pore. To perform MD simulations, the ligand-protein structure after docking was superimposed on the BK_Ca_ channel embedded in the membrane, and the ligand poses were copied. K^+^ and Cl^−^ ions were added to reach a physiological concentration of 0.15 M and obtain the electrically neutralized system. The entire simulation box consisted of approximately 136,600 atoms.

#### 2.6.5. Molecular dynamics (MD) simulations

The MD simulations were performed in periodic boundary conditions to preserve bulk-like behavior and eliminate edge effects. The systems (containing 0, 1, 2, or 4 DBM molecules) were subjected to 10,000 steps of energy minimization using the steepest descent algorithm to remove short contacts that can generate high repulsion energies. Then, the systems were gradually heated from 0 K to 298 K under NVT conditions to maintain a constant volume. At the target temperature of 298 K and under NPT conditions, several equilibration steps were completed before productive MD simulations were started. Firstly, the protein and lipids were kept frozen, and only the water molecules and ions (excluding the four fixed K⁺ ions in the selectivity filter) were allowed to move. This stage ensured proper hydration and charge distribution in the system. Secondly, lipids were released while the protein remained fixed, allowing the membrane to adapt to the protein surface. Then, only the protein backbone atoms were harmonically restrained, while the side chains, lipids, water, and all ions were free to move. Each preliminary MD equilibration step lasted 1 ns. The production runs lasted 200 ns and were performed in three replicates for 8 systems (4 with and 4 without the electric field) in NAMD 3.0 (Humphrey et al., 1996) using the CHARMM36m force field (Huang et al., 2016). Parameters for the ligands were generated in the CGenFF (CHARMM General Force Field) server (Huang et al., 2016). The integration time step was set to 1 femtosecond. The average dimensions of the periodic box were 105±3 Å × 105±3 Å × 123±6 Å. All long-range electrostatic interactions were calculated using the Particle Mesh Ewald (PME) algorithm (Essmann et al., 1998). The van der Waals interactions were smoothly truncated with a switching distance of 10 Å and a cutoff of 12 Å; the pair-list distance was set to 14 Å. Bond lengths involving hydrogen atoms were constrained using the SHAKE algorithm (Kräutler et al., 2001). Temperature control was maintained at 298 K using Langevin thermostat with a damping coefficient of 5 ps⁻¹ applied to all non-hydrogen atoms. Pressure was maintained at 1.01325 bar using a Langevin piston barostat with a period of 50 fs and a decay constant of 25 fs; the group-based pressure coupling was employed. The external electric field was perpendicular to the membrane, allowing transport of potassium ions from the extracellular to the intracellular side of the channel. The electric field strength was set to 10 mV/nm, which corresponds to 40 mV of the membrane potential.

#### 2.6.6. Statistical analysis of pore hydration

The number of water molecules within the defined cylinder encompassing the pore was determined for each frame of the MD trajectories (2000 frames per condition). The cylinder was large enough (determined by the positions of four Cα Tyr318 atoms) to account for the fluctuating pore volume (Fig. 1B). Differences in pore hydration between simulations with different numbers of docked DBM molecules were evaluated using Cohen’s d factor as a standardized measure of effect size. Cohen’s d was calculated as the difference between the mean number of water molecules in two conditions divided by the pooled standard deviation. This metric was chosen to quantify the magnitude of changes independently of sample size, which is large in time-resolved MD simulation data. Effect sizes (d factors) were interpreted according to conventional thresholds (small: d = 0.2, medium: d = 0.5, large: d = 0.8).

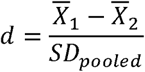

where the pooled standard deviation is a weighted average of standard deviations from two or more independent groups, representing a combined estimate of variability:

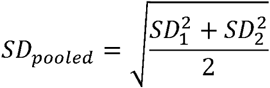

#### 2.6.7. Cavity volume calculation

The pore cavity volume was measured using a cylindrical region of fixed geometry (radius 10 Å, height 12 Å) aligned with the global z axis, which is parallel to the channel’s axis, and dynamically centered as before on the centroid of the four Cα atoms of the Tyr318 pore-lining residue of each subunit for each simulation frame. Volume was estimated using a grid-based method with 2.0 Å spacing. Grid points overlapping protein atoms, defined by their van der Waals radii with an additional 1.0 Å buffer, were excluded. The remaining grid points were multiplied by the grid cell volume to yield the accessible cavity volume (Å³). Water molecules, ions, lipids, and DBM ligands were excluded from the volume calculations. Analyses were performed using MDTraj v1.9.8 and SciPy KDTree v1.16.2.

## 3. Results

### 3.1. DBM is the active component of H. palustris against the BK_Ca_ channel

The activity of BK_Ca_ channels has been previously described in several cell lines, including glioblastoma U87-MG (Wawrzkiewicz-Jałowiecka et al., 2020) and endothelial EA.hy926 cells (Bednarczyk et al., 2013). In these cell lines, the BK_Ca_ channel activity has been observed in both the plasma membrane and mitochondria (Wawrzkiewicz-Jałowiecka et al., 2020). The U87-MG cell line exhibits upregulated BK_Ca_ channel expression (Rosa et al., 2017), making it an excellent model for patch-clamp studies. BK_Ca_ channels display different single-channel conductances (Yang et al., 2015), and consistent with this, BK_Ca_ channels with larger (>280 pS) and smaller (∼160 pS) conductance have been recorded in these cells.

In our study, BK_Ca_ channels recorded from U87-MG and EA.hy926 cells were used to evaluate modulatory effects of purified compounds from *H. palustris* extracts, including dibenzoylmethane, 5-hydroxyflavone, 5-hydroxy-2′-methoxyflavone, and 5-hydroxy-2′,6′-dimethoxyflavone. Among these, only dibenzoylmethane exhibited activity, effectively blocking BK_Ca_ channels in both the intracellular membrane fraction containing mitochondria (Fig. 2A) and the plasma membrane (Fig. 2B). This blocking activity was observed with dibenzoylmethane derived from *H. palustris* (n = 7) as well as chemically DBM (n = 9).

**Fig. 2.**
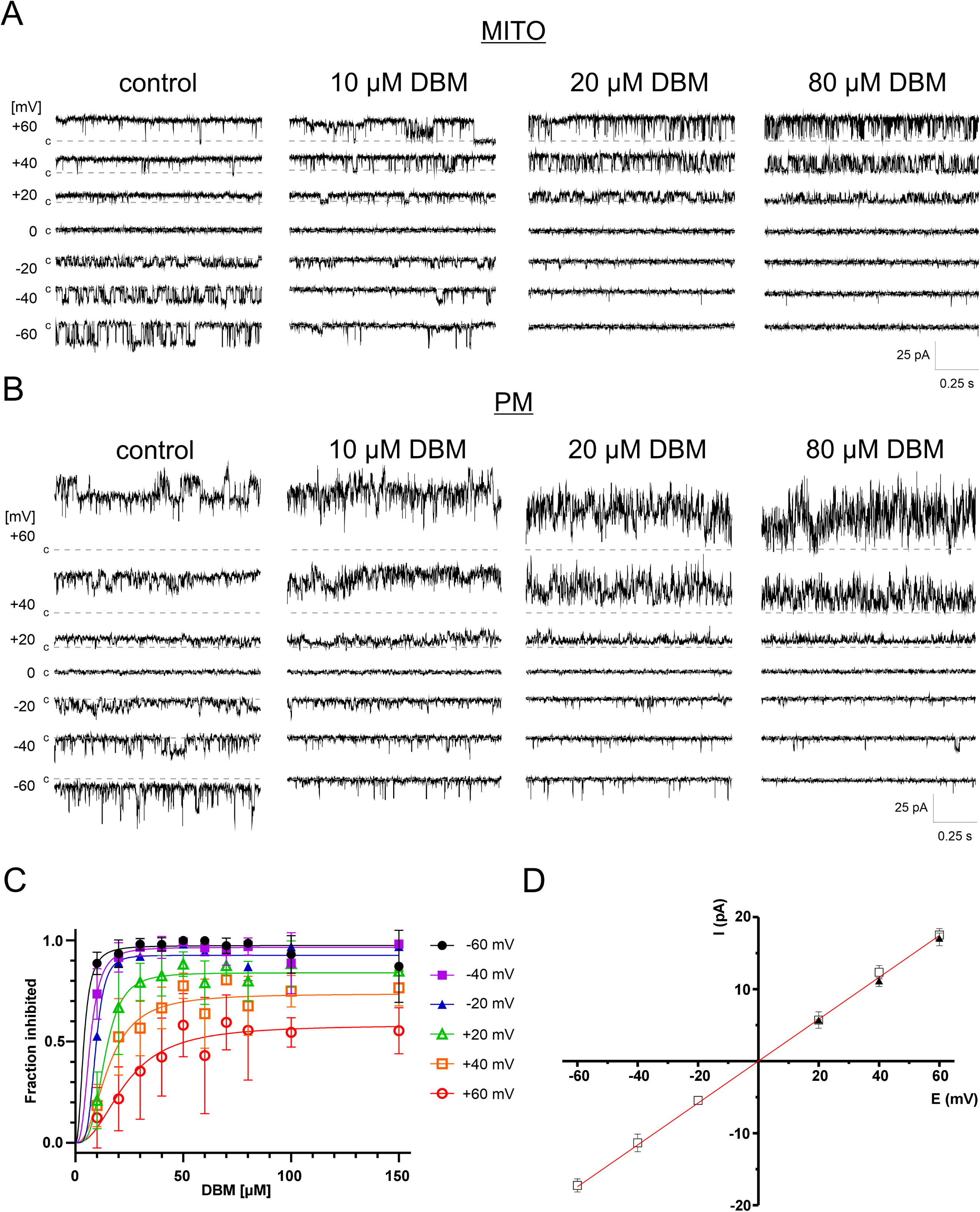
Dibenzoylmethane (DBM) inhibits BK_Ca_ channels in mitochondria and the plasma membrane. (A) Representative single-channel recordings of mitoBK_Ca_ channel activity showing concentration-dependent inhibition by DBM at negative and positive membrane potentials. (B) Representative multi-channel recordings of plasma membrane BK_Ca_ channel activity showing inhibition by DBM at negative and positive membrane potentials. (C) Concentration–response curves showing the voltage dependence of BK_Ca_ channel inhibition by DBM. Data were fitted with the Hill equation (n=2÷6). (D) DBM does not alter the single-channel conductance of BK_Ca_ channels (293 pS). Conductance was estimated using only well-resolved single-channel openings recorded in the presence of 80 µM DBM at positive membrane potentials (n=3). Data were fitted with linear regression.

DBM was a potent BK_Ca_ channel blocker at negative membrane potentials, where BK_Ca_ channels exhibit low open probability (NPo). At -40 mV and -60 mV, more than 80% of channel activity was inhibited with 10 μM DBM (Fig. 2C). However, at positive membrane potentials, DBM acted as a weaker inhibitor. At positive membrane potentials, the concentration dependence of DBM action was best fitted by a saturating Hill-type function. Because of the relatively large variability, the derived parameters should be considered approximate. For +20 mV, the fitted IC_50_ and Hill coefficient were 13.6 µM and 3.6, respectively. For +60 mV, the corresponding values were 23.3 µM and 2.3. By contrast, at negative membrane potentials, the effect was already near maximal at the lowest tested DBM concentrations, consistent with an apparent IC_50_ below 10 µM, and thus did not allow robust estimation of IC_50_ or Hill slope.

Notably, DBM did not completely block channel activity, even at concentrations as high as 200–300 μM, exceeding its calculated solubility limit (logS = − 3.97) and the observed solubility (indicated by solution clouding). This partial inhibition was not due to changes in single-channel conductance (Fig. 2D). Instead, DBM induced a flickering block of BK_Ca_ single-channel currents. At +60 mV, DBM reduced the mean open time from 17.22 ms to 2.74 ms (Fig. 3), leading to a significant decrease in overall NPo.

**Fig. 3.**
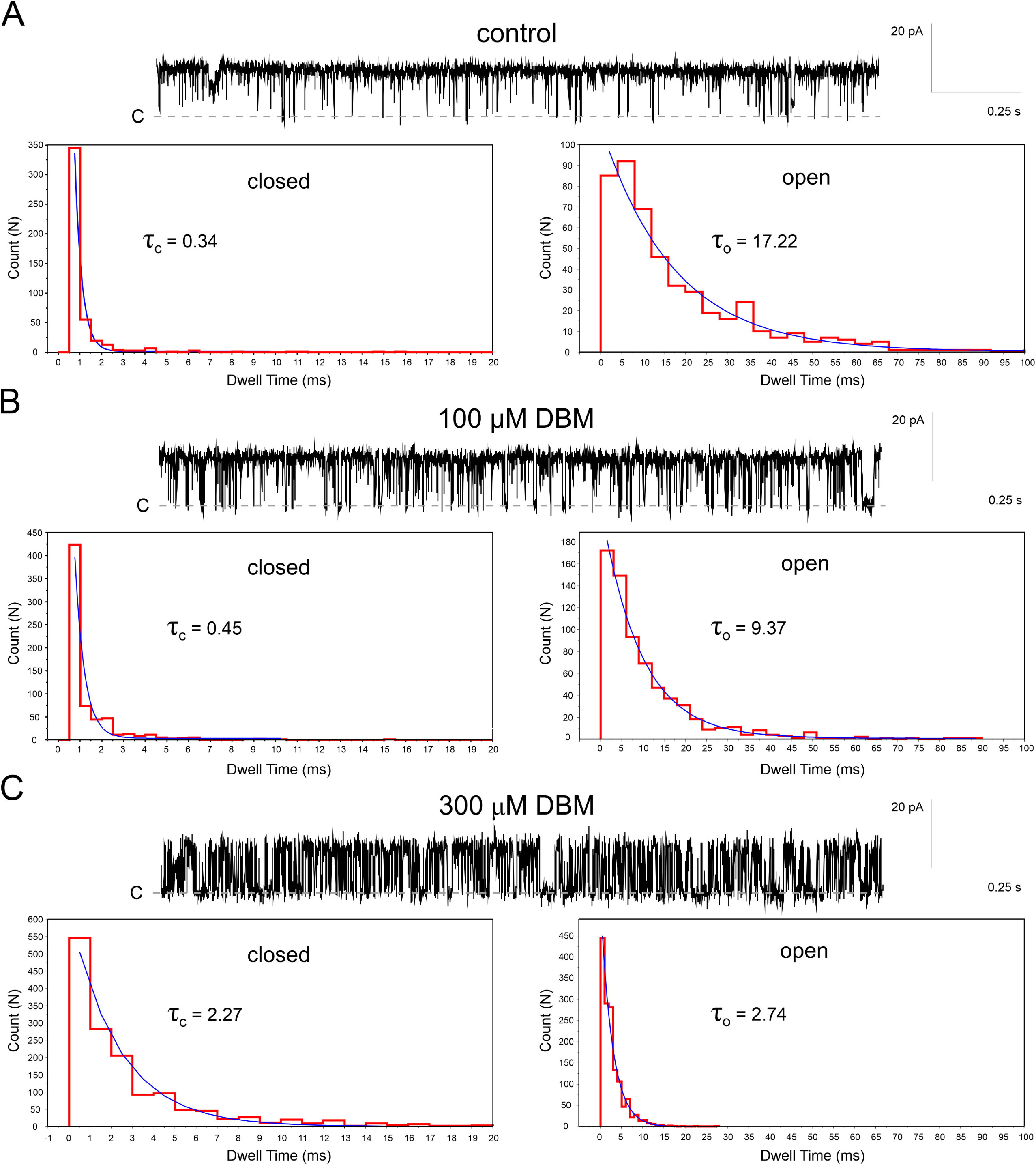
Dibenzoylmethane (DBM) induces a flickering block of BK_Ca_ channels. (A) Representative single-channel recording of BK_Ca_ channel activity under control conditions at +60 mV. Closed- and open-time distributions are fitted by single exponentials with time constants of τ_c_ = 0.34 ms (closed) and τ_o_ = 17.22 ms (open), respectively. (B) Recording in the presence of 100 μM DBM, showing enhanced flickering, with τ_c_ = 0.45 ms (closed) and τ_o_ = 9.37 ms (open). (C) Recording in the presence of 300 μM DBM, showing further enhancement of the flickering block, with τ_c_ = 2.27 ms (closed) and τ_o_ = 2.74 ms (open). Dwell time distributions were calculated from 9 sec recordings.

### 3.2. Dibenzoylmethane binds within the BK_Ca_ channel pore

The above-described behavior suggested that DBM binds with a low affinity within the BK_Ca_ channel pore, obstructing ion flow similarly to the action of quaternary amines (Li and Aldrich, 2004; Tang et al., 2009; Wilkens and Aldrich, 2006). To test this hypothesis, we decided to use a well-known BK_Ca_ channel high-affinity blocker, paxilline (PAX), which binds within the channel pore. PAX binds in the closed channel conformation and traps the channel in the non-conductive state (Zhou and Lingle, 2014). This paxilline block could be overcome by positive membrane potentials, which promote open channel conformation. We hypothesize that, if DBM is a weak open-channel blocker that binds to a site overlapping the paxilline-binding site, it may eventually outcompete PAX when present in molar excess. In the standard protocol used in this work, 0.2 – 2 μM paxilline blocked >90% of BK_Ca_ channel activity (Fig. 4). Next, the paxilline-blocked channels were exposed to up to 100x DBM molar excess. Here, the return of BK_Ca_ channel activity was observed (n=9). This suggests that DBM indeed binds in the channel pore, and its binding site might partially overlap with that of paxilline.

**Fig. 4.**
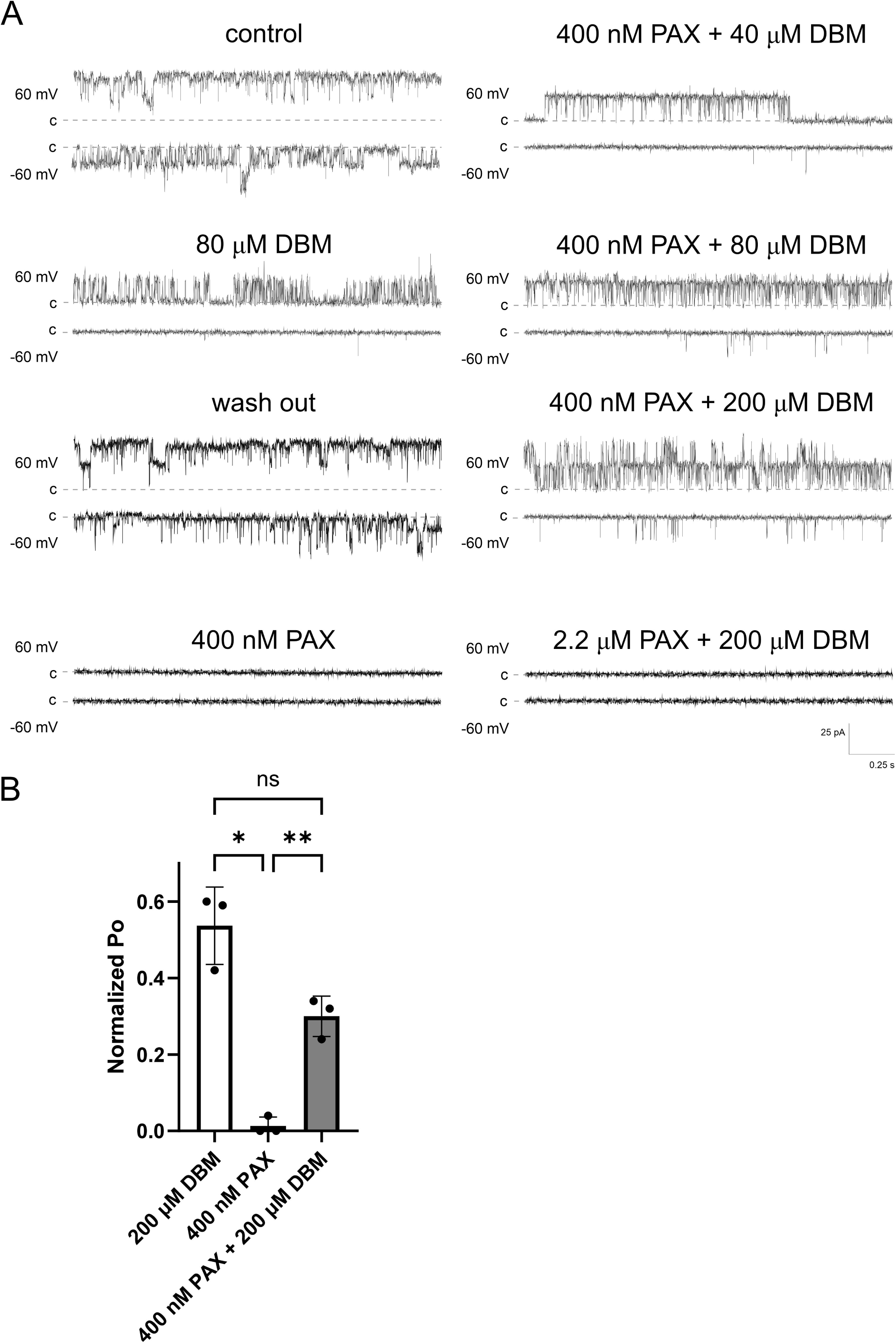
Dibenzoylmethane (DBM) functionally antagonizes paxilline inhibition of BK_Ca_ channels. (A) Representative current traces recorded at ±60 mV under control conditions, after washout, and upon application of 400 nM PAX, 80 μM DBM, and PAX together with increasing concentrations of DBM (40, 80, 200 μM). (B) Normalized open probability (P_o_) of BK_Ca_ channels under the indicated conditions. Excess DBM partially reverses the inhibitory effect of paxilline, consistent with competition for an overlapping or closely related inner-pore binding region. Values are shown as mean ± SD. Welch ANOVA test, p < 0.0032 (n=3).

### 3.3. Diphenyl structure of DBM determines its inhibitory properties

To identify structural determinants critical for DBM binding, we tested three related analogs: 1-phenyl-1,3-butanedione (PBD), trans-chalcone (T-Ch), and (E)-1,3-diphenylprop-2-en-1-ol (DPE) (Fig. 5A).

**Fig 5.**
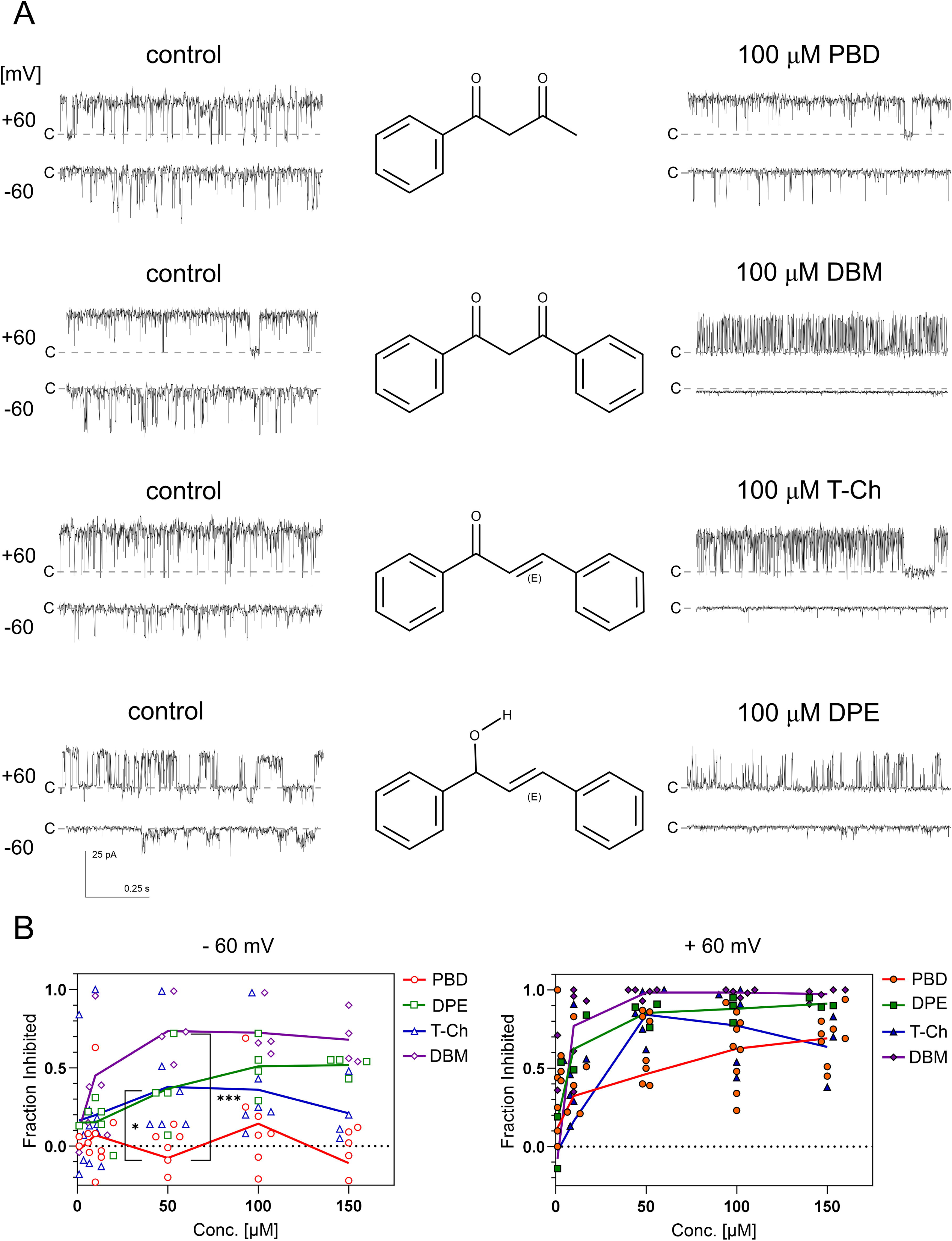
Modulation of BK_Ca_ channel activity by dibenzoylmethane (DBM) and related analogs. (A) Representative single-channel current recordings obtained under control conditions and after application of 100 µM of the indicated compounds: PBD, DBM, T-Ch, or DPE. Traces were recorded at holding potentials of +60 mV and −60 mV. Chemical structures of the tested compounds are shown between the control and drug-treated traces. Dashed lines indicate the closed (c) state. Scale bars: 25 pA, 0.25 s. (B) Concentration–response relationships for inhibition of channel activity at +60 mV (left) and −60 mV (right), expressed as the fraction of activity inhibited relative to control. Individual symbols represent single experiments; lines indicate mean values. PBD is the weakest inhibitor across the tested concentration range. One-way ANOVA test, p < 0.001.

Replacement of one phenyl ring with a methyl group, as in PBD, markedly reduced inhibitory activity, indicating that a diphenyl framework is a key requirement for efficient BK_Ca_ channel block. T-Ch and DPE, both of which preserve two aromatic rings but alter the β-diketone linker, remained active, yet were less effective than DBM, particularly at positive membrane potentials (Fig. 5B). These findings indicate that the diphenyl scaffold is the primary determinant of inhibition, while the β-diketone moiety makes an additional, nonessential contribution that enhances potency or stabilizes the inhibitory binding mode. Thus, the aromatic framework appears to provide the core pore-blocking pharmacophore, whereas the diketo group likely fine-tunes the strength and voltage dependence of the effect.

### 3.4. The presence of DBM results in partial loss of water within the BK_Ca_ channel pore and impaired potassium conductance

To gain mechanistic insight into DBM-mediated inhibition, we employed molecular docking and molecular dynamics (MD) simulations. As a β-diketone, DBM undergoes keto–enol tautomerization and is present in solution as an equilibrium mixture of the diketo form and two equivalent enol tautomers (Fig. 1A). However, due to the flexibility of the diketo form, we chose this tautomer for docking into the inner pore of the BK_Ca_ channel in its open conformation. Notably, the steep concentration dependence of DBM-mediated block (Fig. 2C) suggested that multiple DBM molecules may bind simultaneously within the pore. Consistent with this hypothesis, we were able to dock 1, 2, and 4 DBM ligands in the pore of the BK_Ca_ channel. The docked positions of 1 and 2 ligands were horizontal in relation to the membrane, whereas the positions of 4 ligands were vertical due to a lack of space for horizontal deployment (Fig. 6A).

**Fig. 6.**
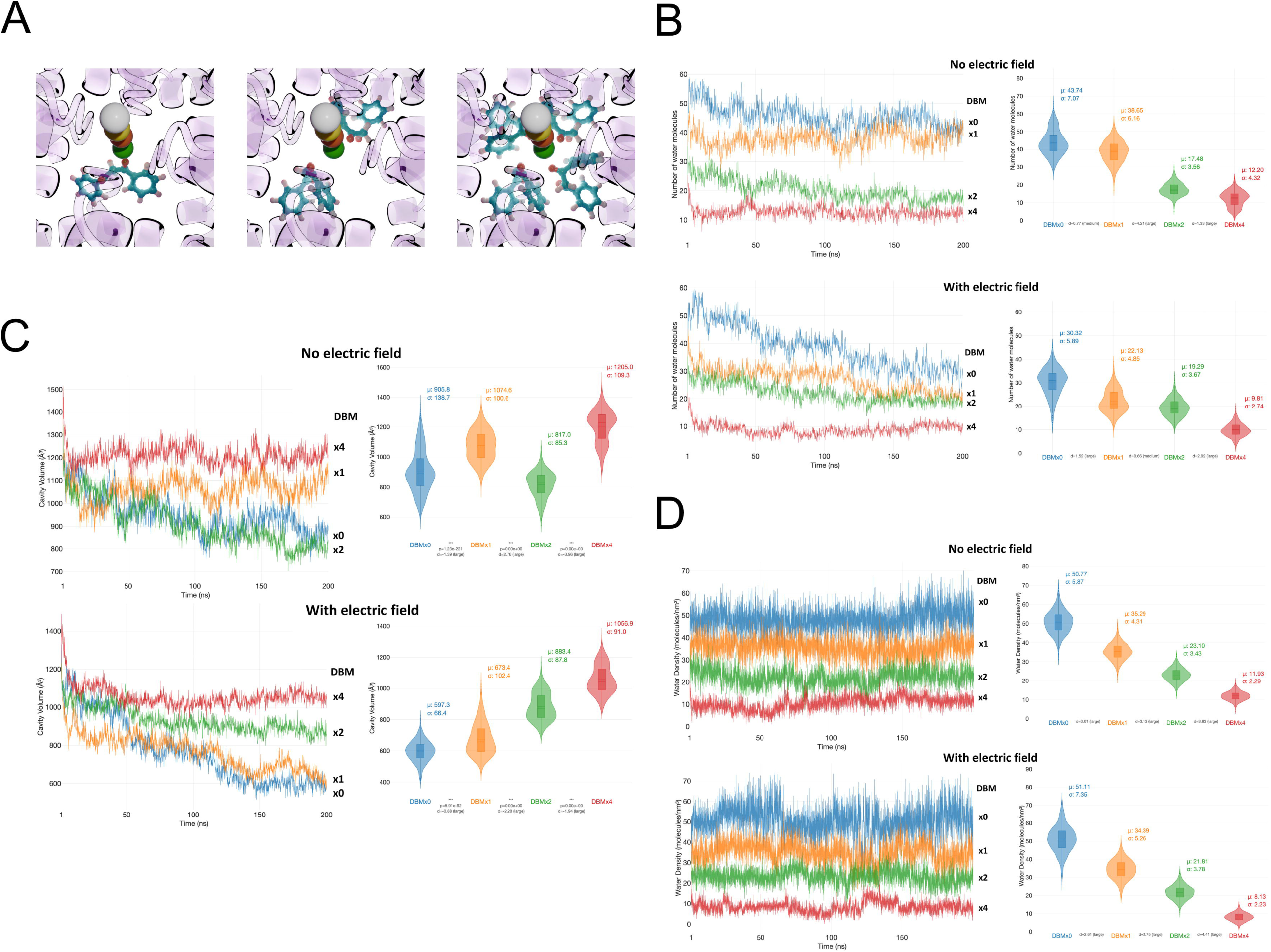
Dibenzoylmethane (DBM) reduces hydration and alters cavity geometry of the BK_Ca_ channel pore. (A) The initial states of the BK_Ca_ channel with 1, 2, and 4 dibenzoylmethane (DBM) ligands in the channel pore. View from the cytoplasmic side. K^+^ ions in the selectivity regions are shown as colored spheres; DBM molecules are shown in ball-and-stick representation. (B) Number of water molecules in the pore during MD simulations of systems containing 0, 1, 2, or 4 DBM molecules, shown as linear plots (left) and violin plots (right). (C) Pore cavity volume during the same simulations, shown as linear plots (left) and violin plots (right). (D) Water density in the pore (number of water molecules divided by pore volume), shown as linear plots (left) and violin plots (right). For panels B–D, data are averaged over three independent simulations per condition. The top row shows simulations without an applied electric field; the bottom row shows simulations with an applied electric field corresponding to a membrane potential of 40 mV. Mean (μ) and standard deviation (σ) values calculated from the final 25% of frames are indicated on the violin plots. Cohen’s d factors are shown between corresponding violin plots. In panel C, p-values were also shown for comparison.

We simulated BK_Ca_ channel systems containing 0, 1, 2, or 4 DBM molecules bound within the pore, embedded in a POPC membrane, and quantified pore hydration (Fig. 6B), the cavity volume (Fig. 6C), and water density (Fig. 6D) under conditions with and without an applied electric field. In MD simulations performed without an electric field, DBM caused a pronounced, copy-number–dependent reduction in the number of water molecules residing in the pore (Fig. 6B). In control simulations lacking DBM (DBMx0), the pore remained highly hydrated, with an average of ∼43 water molecules (Fig. 6B, right upper panel). The presence of a single DBM molecule (DBMx1) resulted in only a modest reduction in hydration (mean ∼38 waters). In contrast, simulations with two or four DBM molecules exhibited a marked loss of pore water, with mean occupancies of ∼17 and ∼12 water molecules, respectively. Notably, both the mean hydration levels and their fluctuations were reduced by approximately half compared to the DBMx0 and DBMx1 conditions, indicating that higher DBM occupancy stabilizes a tightly dehydrated pore environment with limited water exchange.

Application of an electric field further enhanced pore dehydration, particularly in systems with low DBM occupancy (Fig. 6B, lower panels). In the absence of DBM or in the presence of a single DBM molecule, the number of water molecules in the pore decreased substantially over time and exhibited increased temporal variability (Fig. 6B). In control simulations, pore hydration declined from ∼50 water molecules on average in first 25 % of frames to ∼30 on average in last 25 % of frames, suggesting field-driven rearrangements of ions and/or changes in pore geometry. In contrast, systems containing two or four DBM molecules remained persistently dehydrated, with low and relatively stable water occupancies, indicating that DBM dominates pore hydration behavior under these conditions. An interesting case is the system with 1 DBM, since the number of water molecules in the pore is high when no electric field is applied and drops significantly when the electric field is present. Possibly, the behavior of a single DBM molecule in the pore is much different from that in 2 and 4 DBM systems.

Analysis of the pore cavity volume revealed a complementary but distinct dependence on DBM occupancy (Fig. 6C). Under an applied electric field (Fig. 6C lower panels), increasing numbers of DBM molecules progressively increased the mean cavity volume, with DBMx4 exhibiting the largest volumes and DBMx0 the smallest, while DBMx1 and DBMx2 displayed intermediate values. In the absence of an electric field, the DBMx4 system is characterized by the largest volume as before, but then the order is the following: DBMx1 > DBMx0 > DBMx2. The DBMx1 system provided unexpected results in a high number of water molecules (Fig. 6B) and is also in a large pore volume (Fig. 6C), compared to systems with no and 2 DBM molecules.

Unexpected trends found in plots of number of water molecules in the pore and the pore volume can explain the expected trend in water density plots (Fig. 6D). Both for systems without and with the electric field the water density in the pore (number of water molecules divided by the pore volume) the trend is: DBMx0 > DBMx1 > DBMx2 > DBMx4. The average values of water density are stable and do not change upon diminishing of pore volume, which has been very large, especially for the DBMx0 system. Larger σ values for DBMx0 and DBMx1 systems with an electric field, compared to those without an electric field can indicate disturbances in the water structure caused by the passage of ions through the pore. Taken together, these results indicate that DBM exerts dual and decoupled effects on pore structure: it promotes pore dehydration in a strongly copy-number–dependent manner (Fig. 6D) while simultaneously stabilizing the cavity geometry, particularly under an applied electric field. The combination of reduced hydration and physical occupation by DBM is consistent with the formation of a hydrophobic barrier within the pore that is resistant to electric-field–driven narrowing and unfavorable for ion permeation.

We also analyzed movements of ligands (averaged position of geometric center of the ligands) in the pore for BK_Ca_ channels in simulations without and with the electric field (Fig. 7A). The DBM molecules were observed to bind stably within the channel pore throughout the simulations, and although they exhibited movement, they never diffused out of the pore cavity. For the DBMx1 system, the ligand was stably located in the pore center when no electric field was applied, but moved slightly off the center when an electric field was applied. The opposite situation was for the DBMx2 system, where two ligands created an interacting pair and possibly interacted with hydrophobic residues lining the pore. After applying the electric field, the pair of ligands came back to the pore center and interacted with the nearest potassium ion in the selectivity filter, as indicated by the elevated position of the geometric center of the ligands. For the DBMx4 system, nearly the whole pore is full, and its geometry is stabilized by the hydrophobic interactions. After the electric field was applied, the pore became more regular, and the geometric center of ligands aligned with the pore center.

**Fig. 7.**
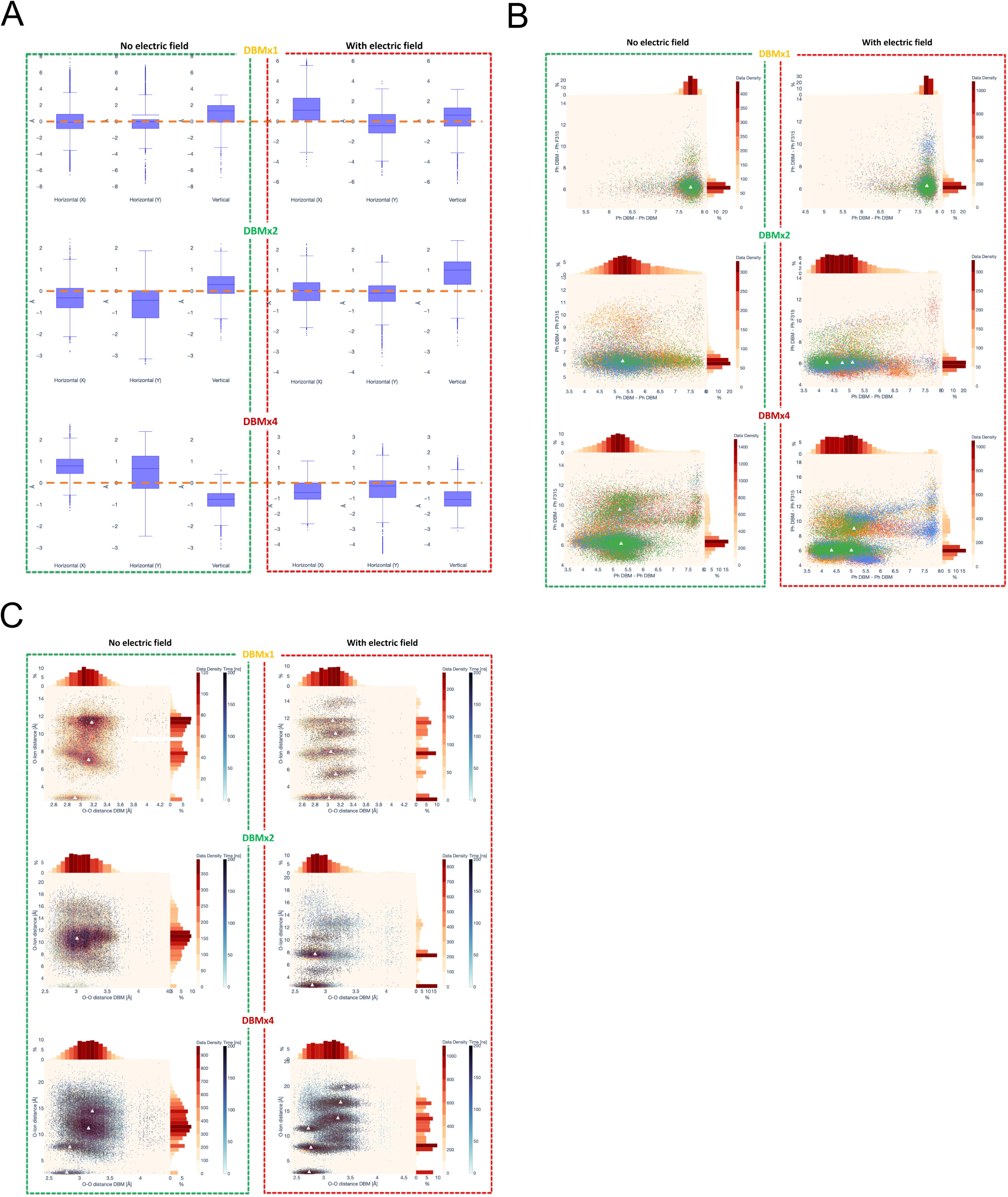
Dynamics and interactions of DBM ligands within the BK_Ca_ channel pore. (A) Positions of DBM ligands (calculated by averaged locations of their geometric centers) in relation to the pore center, which is at (0, 0, 0) coordinates. Each case is averaged over 3 MD simulations without an electric field are shown on the left and simulations with an electric field on the right. (B) 2D scatter plot showing distances between geometric centers of phenyl rings: Ph-Ph among dibenzoylmethane (DBM) molecules vs. Ph DBM-Ph315 averaged over 3 simulations conducted for each system (1, 2, and 4 DBM molecules, without and with the electric field). Major maxima of point density are indicated by white triangles. Affiliation of Tyr315 to the particular subunit is indicated by different colors. (C) 2D scatter plot showing distances involving carbonyl oxygen atoms of dibenzoylmethane (DBM): O-O among DBM molecules vs. O-K^+^ ion averaged over 3 simulations conducted for each system (1, 2, and 4 DBM molecules, without and with the electric field). Major maxima of point density are indicated by white triangles.

Since the DBM ligand is mostly hydrophobic and experiments with DBM analogs indicated the crucial role of phenyl rings, we studied interactions of these rings of the ligands with the side chain of F315 residue, which is in the pore and actively participates in dehydration events (Contreras et al., 2025; Wu et al., 2009; Zhou et al., 2011). DBM ligands can also interact with each other and form π-π interactions. The distance plots for ligand-ligand versus ligand-F315 were created for systems with 1, 2, and 4 ligands in MD simulations without and with the electric field (Fig. 7B). One ligand (DBMx1 system) is far from the F315 residue (6 Å distance to the geometric center of the F315 phenyl ring), and it is in an extended conformation (7.7 Å distance between geometric centers of their phenyl rings). For the DBMx2 system, the distance to Ph-F315 remains the same regardless of subunit, but the Ph-Ph distance between phenyl rings of DBM molecules diminishes to 5.7 Å without an electric field and even further to 4.2 Å with an electric field. Nearly the same situation is for the DBMx4 system, where, after the electric field is applied, the Ph-Ph distance diminishes from 5.2 Å to 4.2 A.

The same type of plots was created for ligand-ligand and ligand-potassium ion distances, but instead of benzene ring centers, the positions of ligand oxygen atoms were taken. The plots were created as before for systems with 1, 2, and 4 (Fig. 7C) ligands in simulations without and with the electric field. For the DBMx1 system, the shortest distance to the K^+^ ion is about 2.5 Å (both without and with the electric field), indicating interaction with the nearest ion in the selectivity filter. The higher-located maxima in the scatter plots indicate distances to higher ions in the selectivity filter. The maxima for MD simulations in the electric field, compared to 2 maxima without an electric field, indicate the strongly organizing role of the electric field on the positions of K^+^ ions. The contour lines for DBM O-O distances are horizontal, revealing the high flexibility of DBM. The same organizing role of the electric field is visible for other DBM systems, with 2 and 4 ligands. For DBMx2, there is nearly no interaction with the nearest K^+^ ion when the electric field is off, but a strong interaction when the electric field is on. For DBMx4, the above interaction is visible for cases without and with the electric field, possible because of the crowded pore and reduced possibility of DBM movements.

To evaluate functional consequences of DBM binding, we analyzed the behavior of DBM and K^+^ ions during 200 ns MD simulations under an applied electric field (Fig. 8). In the exemplary simulation with one DBM (Fig. 8A), the BK_Ca_ channel’s selectivity filter initially contained four K⁺ ions (identical to other MD simulations). First ions passed through the pore very quickly, and a single DBM was not a strong obstacle. Other ions immediately moved to lower positions in the selectivity filter. The second ion exited at about 70 ns, and a third at 120 ns. The last ion in the selectivity filter did not escape till the end of the 200 ns MD simulation. Superimposing all MD simulations (3 simulations for each system) on one graph (Fig. 8B) reveals that among the initial four ions in the selectivity filter, there were 7 passes of K^+^ ions through the pore for the DBMx0 system, 5 for DBMx1, and 4 for DBMx2. There was a complete blockade of ion passing for the DBMx4 system. This indicates increasing difficulty for ions passing the pore with an increasing number of DBM molecules.

**Fig. 8.**
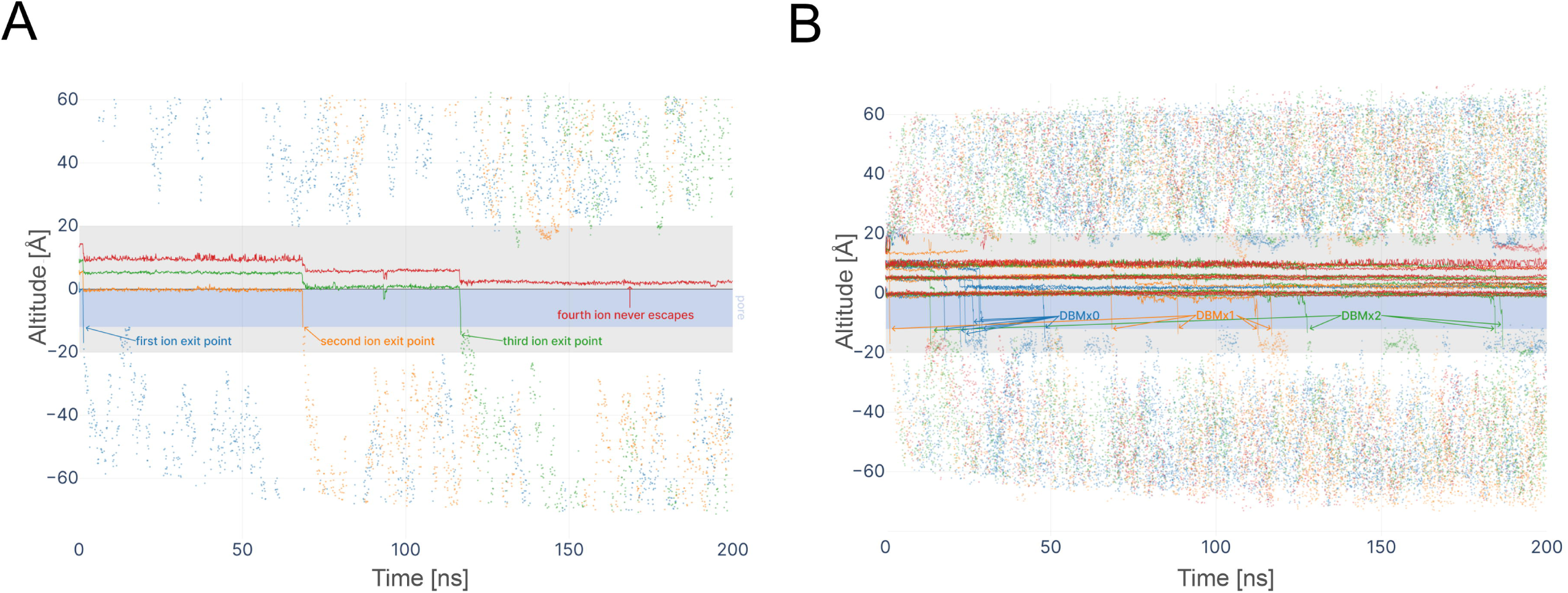
Electric field–driven K⁺ permeation through the BK_Ca_ pore in the presence of dibenzoylmethane (DBM). Time evolution of the axial position (altitude along the membrane normal) of individual K⁺ ions during MD simulations with an applied electric field. Continuous traces represent ion trajectories within the selectivity filter (above 0 level) and the channel pore (below 0 level within the blue area indicating the pore). The membrane spans −20 Å to +20 Å and is shown in grey. The scattered points indicate ion positions outside the channel. Only trajectories of four K^+^ ions initially located in the selectivity filter are shown. (A) Representative MD simulation showing trajectories of ions in the BK channel with one DBM molecule present in the pore. Colors indicate different K^+^ ions. (B)) Superposition of trajectories from all simulations (three simulations per condition) for systems containing 0, 1, 2, or 4 DBM molecules. Colors indicate the different systems. No K⁺ escape was observed in simulations containing four DBM molecules.

In simulations containing DBM, the motion of K⁺ ions in the selectivity filter was markedly altered relative to the control (Movie 1). With 1 DBM present (Movie 2), the filter could still initially host up to three K⁺ ions, but their dynamics were constrained. K⁺ ions remained in the filter longer and did not progress outward in the usual rapid succession observed in a knock-on event. K⁺ ions appeared to become “trapped” in the filter or just below it. Here, the DBM molecule remained lodged in the central cavity for the duration of the simulation and it adopted a stable orientation – its two aromatic rings nestled against hydrophobic surfaces of the pore-lining helices, and its β-diketone moiety oriented toward the toward K^+^ in the selectivity filter for most of the time– suggesting favorable interaction with that adjacent K⁺ ion, which observed in all simulations.

In the simulation with two DBMs (Movie 3), K⁺ permeation was severely impaired: the ions in the filter mostly oscillated in place or exhibited only local fluctuations, with only one exiting to the intracellular side after a significant delay. Both DBM molecules found persistent binding sites in the pore region, often adopted a parallel configuration, with their keto groups directed toward the K^+^ ion in the conduction pathway. These two ligands remained relatively stable, occasionally wobbling or reorienting, but largely stayed in proximity to the conduction pathway.

Introducing four DBMs eliminated any observable K⁺ conduction (Movie 4). With four molecules present, the channel pore became crowded with inhibitors. Multiple DBMs clustered within the central cavity – forming a collective physical barrier. Interestingly, two DBMs adopted paralel configuration with their keto groups oriented towards the K^+^ ion, while the additional two DBMs displaced toward the lower part of the cavity in a less ordered fashion. All four DBM molecules remained bound around the pore region (none drifted far into the bulk solution), indicating a strong tendency for DBM to partition into and stay within the channel’s inner pore/vestibule.

The overall stability of DBM binding was high in all cases, implying that once a DBM molecule occupied the pore, it tended not to escape it on the timescale of the simulations.

## 4. Discussion

Our study identified dibenzoylmethane (DBM), a β-diketone derived from *Hottonia palustris*, as a novel pore blocker of BK_Ca_ channels. DBM was found in other plants, i.e., licorice (*Glycyrrhiza glabra*) (Mancia et al., 2014) or feijoa (*Acca sellowiana*) (Peng et al., 2019). Interestingly, DBM is also generated as a byproduct of thermal- and photo-oxidative degradation of polystyrene in the presence of sand or silica (Costa et al., 2024). Given the global prevalence of plastic pollution, such abiotic degradation processes may be a significant environmental source of DBM. Functionally, DBM exhibits pleiotropic biological activities, many of which may be mediated, at least in part, through inhibition of BK_Ca_ channel function. Among its most extensively studied effects are anticancer actions. DBM induces cell cycle arrest (Jackson et al., 2002), suppresses oncogenic signaling pathways, and attenuates the proliferation of prostate cancer cells by inhibiting androgen receptor expression (Jackson et al., 2007; Khor et al., 2009). In breast carcinoma cells (MCF-7), DBM was shown to inhibit phorbol ester-induced migration and invasion (Liao et al., 2015).

Beyond its antineoplastic effects, DBM also exerts anti-inflammatory and antioxidant actions. It significantly reduces lipopolysaccharide (LPS)-induced nitric oxide (NO) production by downregulating inducible nitric oxide synthase (iNOS), as well as pro-inflammatory cytokines such as TNF-α, IL-1β, and IL-6 (Kang et al., 2018). Moreover, DBM protects against inflammation-associated oxidative stress, including the death of cultured neuronal cells and obesity-related chronic inflammation (You and Choi, 2023).

DBM has also been reported to confer metabolic benefits. In mice fed a high-fat diet, DBM attenuated weight gain and prevented hepatic and abdominal fat accumulation (Kim et al., 2015). In skeletal muscle cell lines, it promoted AMP-activated protein kinase (AMPK) phosphorylation and enhanced glucose uptake. Additionally, in preadipocytes, DBM suppressed the activity of acetyl-CoA carboxylase, a key enzyme in lipogenesis (Kim et al., 2015). Beneficial effects of co-treatment of DBM with diazoxide (DZ), a potassium channel activator, on spatial memory deficits and buildup of hippocampal Aβ plaques in Alzheimer’s disease rat model were observed (Wallace et al., 2024).

Taken together, these findings suggest that DBM influences multiple cellular pathways associated with proliferation, inflammation, metabolism, and redox regulation. Given that BK_Ca_ channels are sensitive to intracellular Ca²⁺, redox state, and metabolic signals, the broad pharmacological profile of DBM is consistent with its potential role as a modulator of BK_Ca_ channel activity.

Our electrophysiological recordings demonstrated that DBM inhibits BK_Ca_ channels in a dose- and voltage-dependent manner. The steep inhibition-concentration relationship, along with molecular docking and MD simulations, suggests that multiple DBM molecules can simultaneously occupy the channel pore, consistent with a cooperative or multivalent binding mechanism.

In single-channel recordings, DBM induced a characteristic flickering block, evidenced by decreased mean open times and increased closed times. This pattern is typical of open-channel blockers that transiently obstruct the ion conduction pathway, such as quaternary ammonium ions (Li and Aldrich, 2004). Importantly, DBM was also able to outcompete paxilline, a high-affinity pore blocker of BK_Ca_ channels. Paxilline is known to stabilize the closed state of the channel, preventing ion conduction by promoting a non-conductive conformation of the gate (Zhou et al., 2020). In contrast, DBM appears to bind to the open state, as suggested by the partial reversal of paxilline block when excess DBM was applied. This competitive interaction implies that DBM and paxilline share overlapping binding regions within the pore, despite potentially targeting different conformational states. On the other hand, DBM affects the BK_Ca_ channel more at negative membrane potential, suggesting that concerted behavior with the closed channel.

The ability of DBM to compete with paxilline despite its lower affinity highlights its potential utility as a molecular scaffold for the development of novel BK_Ca_ inhibitors. DBM structure, like that of paxilline and other pore-targeting BK_Ca_ modulators such as quercetin (Kampa et al., 2022), contains aromatic rings, which may facilitate π–π stacking and hydrophobic interactions within the pore environment including hydrophobic gating discussed in further paragrah.

The structure–activity data obtained with DBM analogs further refine this mechanistic interpretation. Replacement of one phenyl ring with a methyl group, as in PBD, markedly reduced inhibitory efficacy, indicating that preservation of the diphenyl scaffold is a key requirement for effective BK_Ca_ channel block. By contrast, T-Ch and DPE, both of which retain two aromatic rings but alter the β-diketone linker, remained inhibitory, yet their effects were weaker than those of DBM, particularly at positive membrane potentials. This pattern indicates that the diphenyl aromatic surface provides the principal hydrophobic framework required for productive pore occupancy, whereas a monoaryl analog such as PBD may be too small to achieve comparable occupancy or to efficiently promote local dewetting of the conduction pathway. In addition, the β-diketone moiety is not absolutely required for activity but enhances inhibitory efficacy. Mechanistically, the two phenyl rings likely promote stable partitioning into the inner cavity and favor ligand–ligand as well as ligand–pore hydrophobic and π-stacking interactions, while the diketo group may further optimize ligand orientation and strengthen local polar interactions that stabilize the inhibitory binding mode. Thus, the diphenyl scaffold appears to constitute the core pharmacophoric element, whereas the chemical nature of the linker between the aromatic rings modulates the strength and voltage dependence of channel inhibition.

Experimental studies on aromatic β-diketones, including DBM derivatives, demonstrate that keto–enol tautomerism is a prominent feature of this scaffold and can be shifted by chemical context (Zawadiak and Mrzyczek, 2012). In the present work, DBM was modeled in the diketo form during docking for methodological rather than mechanistic reasons. Most mainstream docking workflows employ fixed-topology, fixed-protonation ligand models and do not explicitly simulate proton transfer or tautomeric interconversion during sampling (Martin, 2009). Under these constraints, ligand tautomer assignment is a necessary preprocessing step, and alternative protomer/tautomer choices can measurably affect predicted poses and rankings (Brink and Exner, 2009). Our primary objective was to use docking as a geometric and hypothesis-generating tool to identify sterically plausible binding regions and orientations within the BK_Ca_ channel pore, rather than to claim an absolute binding free energy or definitive bound tautomer. This framing is consistent with published guidance that docking is most reliable for generating interaction hypotheses and prioritizing poses for subsequent validation, while scoring functions and rank ordering can be unreliable for quantitative affinity prediction (Bender et al., 2021; Ramírez and Caballero, 2016). In this context, use of the diketo form provides a conservative and reproducible representation of DBM that enables consistent sampling across runs and straightforward comparison of candidate poses. Importantly, the docking results obtained with the diketo form are interpreted here as structural hypotheses (putative orientations and contact regions) rather than evidence that DBM binds the BK_Ca_ pore predominantly as the diketo tautomer under physiological conditions.

A key mechanistic insight from our study is that DBM exploits the hydrophobic gating mechanism of the BK_Ca_ channel. In hydrophobic gating, a sufficiently nonpolar region within the pore can spontaneously de-wet, creating an energy barrier to ion and water entry even when the pore is geometrically open (Aryal et al., 2015; Yazdani et al., 2020). Previous structural and computational studies have shown that BK_Ca_ channel activity can be regulated by changes in pore hydration, independently of canonical gating conformations (Coronel et al., 2025; Jia et al., 2018; Nordquist et al., 2023). MD simulations revealed that DBM induces partial dehydration of the pore, a consequence of its hydrophobic phenyl groups displacing water molecules from the channel interior. This reduction in pore water occupancy, especially around the central cavity and the lower selectivity filter, elevates the energetic barrier for ion movement. Since water molecules are essential for coordinating and stabilizing K⁺ ions during permeation, DBM-induced dehydration functionally “locks” the channel in a non-conductive state, without requiring closure of the activation gate.

In this context, DBM appears to act as a hydrophobic inverse of known BK_Ca_ openers such as NS11021, which promotes conduction by increasing pore hydration (Nordquist et al., 2024; Rockman et al., 2020). DBM does the opposite: by inserting into the hydrophobic inner cavity, it reduces water occupancy and stabilizes a dry, non-conductive pore environment.

The inhibition of BK_Ca_ channels by DBM in both mitochondrial and plasma membrane fractions suggests broad applicability in physiological contexts where these channels play critical roles. For instance, mitochondrial BK_Ca_ channels are implicated in cellular bioenergetics and protection against oxidative stress (Szewczyk, 2024). Targeting these channels with DBM or its derivatives could provide therapeutic benefits in conditions such as ischemia-reperfusion injury or neurodegenerative diseases. However, the weak blocking efficacy of DBM at positive membrane potentials may limit its effectiveness under certain physiological conditions. Future studies should focus on optimizing DBM’s pharmacological properties, such as its solubility and binding affinity.

While this study provides insights into DBM’s interaction with BK_Ca_ channels (both in plasma membrane and the mitochondrial membrane), several limitations must be addressed. First, the exact structural determinants of DBM’s binding site within the pore remain unclear and require high-resolution structural studies. Second, the low water solubility of DBM presents a challenge for *in vivo* applications and necessitates the development of more soluble derivatives. Finally, the physiological relevance of DBM’s pore-blocking activity should be validated in more complex systems, such as primary cells or animal models. DBM was studied in the context of its biological effects. Future research should aim to elucidate the precise molecular interactions between DBM and BK_Ca_ channels, explore structure-activity relationships, and assess the therapeutic potential of DBM-based compounds. Combining experimental approaches with computational modeling will be essential for advancing our understanding of this promising new class of BK_Ca_ channel blockers.

## 5. Conclusion

Our results identify dibenzoylmethane (DBM) as a novel pore-directed inhibitor of BK_Ca_ channels in mitochondrial and plasma membranes. DBM reduces channel activity by inducing a flickering block consistent with transient occupancy of the inner pore, without altering single-channel conductance. Docking and MD simulations support this mechanism and indicate that DBM favors a less hydrated, non-conductive pore state. Together with the analog data, these findings indicate that DBM is a useful scaffold for studies and future optimization of pore-targeting BK_Ca_ modulators.

## CRediT authorship contribution statement

**Piotr Koprowski:** Conceptualization, Methodology, Validation, Formal analysis, Investigation, Resources, Data curation, Writing-original draft, Visualization. **Przemysław Miszta:** Conceptualization, Data curation, Investigation, Formal analysis, Methodology, Software, Visualization. **Jakub W. Strawa:** Resources.**Yurii Krempovych:** Software, Methodology, Visualization. **Alicja Ziajowska:** Investigation, Software. **Sławomir Filipek:** Methodology, Writing-review&editing, Visualization. **Adam Szewczyk:** Conceptualization, Funding acquisition, Project administration, Supervision, Writing-review&editing. **Michał Tomczyk:** Conceptualization, Resources, Writing-review&editing.

## Acknowledgments

We would like to thank Prof. Piotr Bednarczyk from the Department of Physics and Biophysics, Warsaw University of Life Sciences, for helpful discussions

## Funding

This study was supported by the Polish National Science Centre [grant no. 2023/51/B/NZ5/00999] to A.S.

## Declaration of competing interest

The authors declare that they have no known competing financial interests or personal relationships that could have appeared to influence the work reported in this paper.

## Data availability

Data will be made available on request.

## Notes

### Competing Interest Statement

The authors have declared no competing interest.

